# Single-cell RNA counting at allele- and isoform-resolution using Smart-seq3

**DOI:** 10.1101/817924

**Authors:** Michael Hagemann-Jensen, Christoph Ziegenhain, Ping Chen, Daniel Ramsköld, Gert-Jan Hendriks, Anton J.M. Larsson, Omid R. Faridani, Rickard Sandberg

## Abstract

Large-scale sequencing of RNAs from individual cells can reveal patterns of gene, isoform and allelic expression across cell types and states^1^. However, current single-cell RNA-sequencing (scRNA-seq) methods have limited ability to count RNAs at allele- and isoform resolution, and long-read sequencing techniques lack the depth required for large-scale applications across cells^2,3^. Here, we introduce Smart-seq3 that combines full-length transcriptome coverage with a 5’ unique molecular identifier (UMI) RNA counting strategy that enabled *in silico* reconstruction of thousands of RNA molecules per cell. Importantly, a large portion of counted and reconstructed RNA molecules could be directly assigned to specific isoforms and allelic origin, and we identified significant transcript isoform regulation in mouse strains and human cell types. Moreover, Smart-seq3 showed a dramatic increase in sensitivity and typically detected thousands more genes per cell than Smart-seq2. Altogether, we developed a short-read sequencing strategy for single-cell RNA counting at isoform and allele-resolution applicable to large-scale characterization of cell types and states across tissues and organisms.

---

Most scRNA-seq methods count RNAs by sequencing a UMI together with a short part of the RNA (from either the 5’ or 3’ end)^4^. These RNA end-counting strategies have been effective in estimating gene expression across large numbers of cells, while controlling for PCR amplification biases, yet RNA-end sequencing has seldom provided information on transcript isoform expression or transcribed genetic variation. Moreover, many massively parallel methods suffer from rather low sensitivity (i.e. capturing only a low fraction of RNAs present in cells)^5^. In contrast, Smart-seq2 has combined higher sensitivity and full-length coverage^6^, which e.g. enabled allele-resolved expression analyses^7^, however at a lower throughput, higher cost and without the incorporation of UMIs. Sequencing of full-length transcripts using long-read sequencing technologies could directly quantify allele and isoform level expression, yet their current depths hinder their broad application across cells, tissue and organisms^2,3^. To overcome these shortcomings, we sought to develop a sensitive short-read sequencing method that would extend the RNA counting paradigm to directly assign individual RNA molecules to isoforms and allelic origin in single cells.

## Results

We systematically evaluated reverse transcriptases and reaction conditions that could improve the sensitivity, i.e. the number of RNA molecules detected per cell, compared to Smart-seq2^6^. Our efforts were focused on improving a Smart-seq2 like assay that retains full-length transcript coverage, thus consisting of oligo-dT priming, reverse transcription followed by template switching, full cDNA amplification using PCR and finally Tn5-based tagmentation and library construction (**Figure 1a**). After assessing hundreds of different reaction conditions in HEK293T cells, with the 96 most notable conditions sequenced (**Figure S1** and **Table S1**), the highest sensitivity was obtained using Maxima H-minus reverse transcriptase (hereafter called Maxima), in line with recent work^8^. We noted that switching the salt during reverse transcription from KCl to NaCl or CsCl improved sensitivity in Maxima-based single-cell reactions compared to standard KCl conditions (**Figure S2**), likely due to reduced RNA secondary structures^9^. Moreover, performing reverse transcription in 5% PEG improved yields, as recently demonstrated^8^, and we added GTPs^10^ or dCTPs to stabilize or promote the template switching reaction (**Figure S2**). We tested a number of DNA polymerase enzymes, however KAPA HiFi Hot-Start polymerase remained most compatible with the reaction chemistry and yielded highest sensitivity. Importantly, we constructed a template-switching oligo (TSO) that harbored a primer site consisting of a partial Tn5 motif^11^ and a novel 11 bp tag sequence, followed by a 8bp UMI sequence and three riboguanosines, the latter hybridizes to the non-templated nucleotide overhang at the end of the single-stranded cDNA. After sequencing, the 11 bp tag can be used to unambiguously distinguish 5’ UMI-containing reads from internal reads (**Figure 1a**). Therefore, we obtain strand-specific 5’ UMI-containing reads and unstranded internal reads spanning the full-transcript without UMIs in the same sequencing reaction (**Figure 1b**). The proportions of 5’ to internal reads could be tuned by altering the Tn5-based tagmentation reaction (**Figure 1c**). We termed the final protocol Smart-seq3, and it significantly improved the detection of polyA+ protein-coding (**Figure 1d**) and non-coding RNAs (**Figure S3**) in HEK293FT cells. Compare to Smart-seq2, the cell-to-cell correlations in gene expression profiles improved significantly with Smart-seq3 (**Figure 1e**) and we uncovered remarkable complexity in the HEK293T cell transcriptomes with up to 150,000 unique molecules detected (**Figure 1f**). Strikingly, comparison of Smart-seq3 to single-molecule RNA-FISH revealed that Smart-seq3 detected up to 80% of the molecules detected by smRNA-FISH per cell^12^, and on average 69% of smRNA-FISH molecules across the four genes tested (**Figure 1g,h**). Altogether, this demonstrated that Smart-seq3 has significantly increased sensitivity compared to Smart-seq2 and is even approaching the sensitivity of smRNA-FISH.

**Figure 1.**
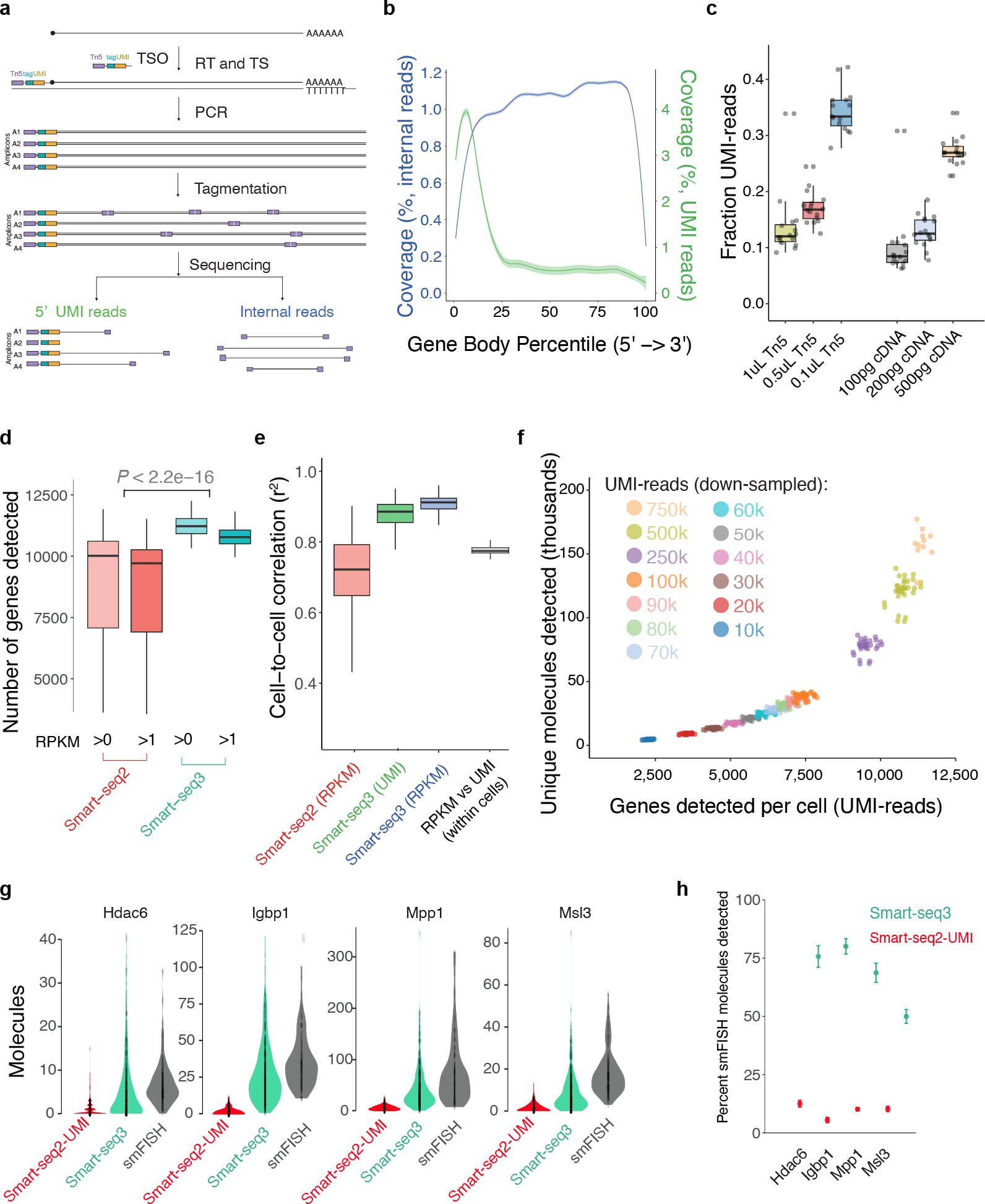
Overview of single-cell RNA-sequencing in Smart-seq3. **(a)** Library strategy for Smart-seq3. PolyA+ RNA molecules are reverse transcribed and template switching is carried out at the 5’ end. After PCR preamplification, tagmentation via Tn5 introduces near-random cuts in the cDNA, producing 5’ UMI-tagged fragments and internal fragments spanning the whole gene body. (**b**) Gene body coverage averaged over HEK293FT (n = 96) cells sequenced with the Smart-seq3 protocol. Shown is the mean coverage of UMI reads (green) and internal reads (blue) shaded by the standard deviation. (**c**) Effect of tagmentation conditions on the fraction of UMI-containing reads (16 HEK293FT cells per condition). Left panel: varying Tn5 with constant 200 pg cDNA input. Right panel: varying cDNA input with constant 0.5ul Tn5. (**d**) Gene detection sensitivity for Smart-seq2 (44 cells) and Smart-seq3 (88 cells), downsampled to 1 million raw reads per HEK293FT cell. Shown are number of genes detected over 0 or 1 RPKM. P-value was computed as a two-sided *t*-test. (**e**) Reproducibility in gene expression quantification across HEKF293FT cells for Smart-seq2 (44 cells) and Smart-seq3 (88 cells) at RPKM and UMI level. Shown are adjusted r^2 for all pairwise cell to cell linear model fits in libraries downsampled to 1 million reads per cell. (**f**) Sensitivity to detect RNA molecules in Smart-seq3 shown by summarizing the number of unique error-corrected UMI sequences and genes detected per HEK293FT cell. Colors indicate the per cell downsampling depth ranging from 10.000 (n = 24 cells) to 750.000 (n = 16 cells) UMI-containing sequencing reads. (**g**) Violin plots summarizing the number of molecules detected per cell with Smart-seq2-UMI, Smart-seq3 and using smRNA-FISH for four X chromosomal genes (Hdac6, Igbp1, Mpp1 and Msl3). (**h**) Estimating the percent of smRNA-FISH molecules that were detected in cells using Smart-seq2-UMI and Smart-seq3. Shown are means and 95% confidence intervals.

We next developed a strategy for the *in silico* reconstruction of RNA molecules. Importantly, the PCR preamplification of full-length cDNA in Smart-seq3 is followed by Tn5 tagmentation, so copies of the same cDNA molecule with the same UMI obtain variable 3’ ends that map to different parts of the specific transcript (**Figure 2a**). Therefore, paired-end sequencing of these libraries results in 3’ end sequences that span different parts of the initial cDNA molecule that we computationally can link to the specific molecule based on the 5’ UMI sequence, thus enabling parallel reconstruction of the RNA molecules (**Figure 2a**). To experimentally investigate the RNA molecule reconstructions, we created Smart-seq3 libraries from 369 individual primary mouse fibroblasts (F1 offspring from CAST/EiJ and C57/Bl6J strains) that we subjected to paired-end sequencing. Aligned and UMI-error corrected read pairs^13^ were investigated and linked to molecules by their UMI and alignment start coordinates. An example of read pairs that were derived from a particular molecule transcribed from the *Cox7a2l* locus in a single fibroblast is visualized in **Figure S4**. We then explored how often the reconstructed parts of the RNA molecules covered strain-specific single-nucleotide polymorphisms (SNPs). Strikingly, unambiguous identification of allelic origin by direct sequencing of SNPs in reads linked to the UMI was observed for 61% of all detected molecules (**Figure 2b**), with increasing assignment percentage with increasing SNP density within transcripts (**Figure 2c**). Previous single-cell studies estimated allelic expression as the product of the RNA quantification (in molecules or RPKMs) and fraction SNP-containing reads supporting each allele^7,12,14^, and we next investigated how those estimates compared to the direct allelic RNA counting made possible with Smart-seq3. Reassuringly, allelic expression estimates and direct allelic RNA counting showed good overall correlation when aggregated over cells (**Figure 2d**). Moreover, using a linear model to quantify the agreement of the two measures across genes within cells revealed a strong correlation (Spearman rho=0.82±0.08 and slope=0.88±0.06) without any apparent bias (intercept=0.06±0.03) (**Figure 2e**). Thus, direct allelic RNA counting is feasible in single cells and validates previous efforts to estimate allelic expression from separated expression and allelic estimates in single cells^7,12,14^.

**Figure 2.**
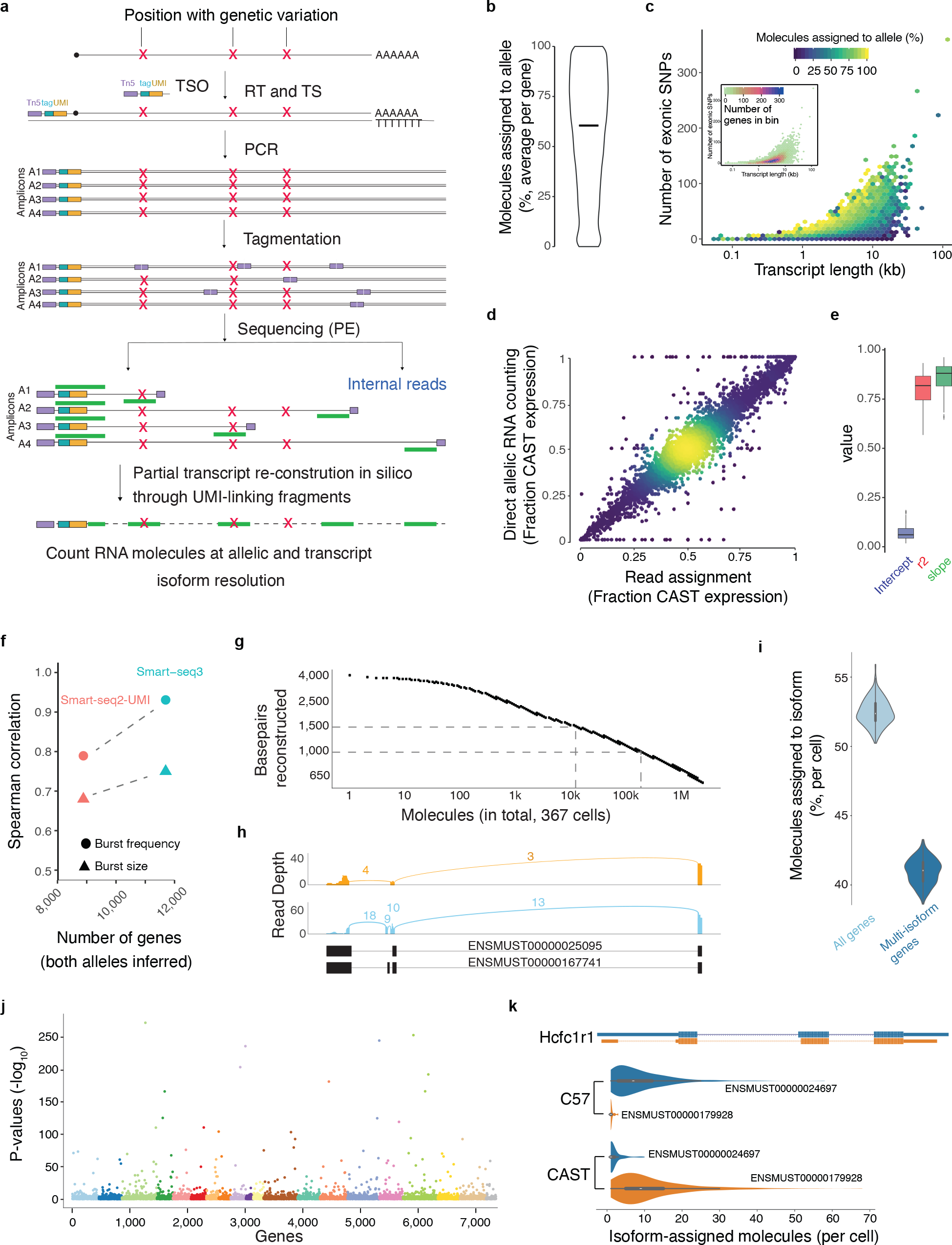
Single-cell RNA counting at allele and Isoform-resolution. **(a)** Strategy for obtaining allelic and isoform resolved information using Smart-seq3. Red crosses indicate transcript positions with genetic variation between alleles. After tagmentation, UMI fragments are subjected to paired-end sequencing (indicated in green), linking molecule-counting 5’ ends with various gene-body fragments that can cover allele-informative variant positions and spanning isoform-informative splice junctions, thus allowing in silico reconstruction of isoforms and allele of origin. **(b)** Average percentage of molecules that could be assigned to allele origin based on covered SNPs, from 369 individual CAST/EiJ × C57/Bl6J hybrid mouse fibroblasts. Only genes detected in >5 % of cells were considered (n = 15,158 genes). **(c)** Effect of transcript length and number of exonic SNPs on allele assignment of RNA molecules. Shown are genes (n = 15,158) grouped into 50 2D-bins colored by the average gene-wise percentage of molecules assigned to allele of origin. Inset shows the number of genes per visualized bin. **(d)** Concordance of allele expression from RNA counting and traditional estimates based on separated expression and allele-fractions from internal reads. Shown are the average CAST allele fractions for 15,158 genes over 369 mouse Fibroblasts. Dots are colored by the local density of data points. **(e)** Results from linear models that compared direct allelic RNA counting with previous read-based estimates of allelic expression, within each of 369 individual fibroblasts. For each cell (n = 369), we computed a linear model fit of CAST allele fraction between direct reconstructed molecule assignment and traditional read-based estimates. Shown are boxplots of the Intercept, slope and r^2 values obtained from each linear model per cell. **(f)** Demonstrating the improved abilities of Smart-seq3 to infer transcriptional burst kinetics compared to Smart-seq2-UMI (the Smart-seq2 chemistry combined with a UMI in the TSO). Inference was made in F1 CAST/EiJ × C57/Bl6J mouse fibroblasts and we show the spearman correlation between the CAST and C57 kinetics across genes for burst size and frequency. Additionally, the x-axis shows the number of genes for which we could reliably infer the bursting kinetics. **(g)** Summarizing the numbers of RNA molecules (x-axis, log10) reconstructed to different lengths (in base pairs, y-axis), showing only molecules additionally assigned to a unique transcript isoform. In total, the one million longest reconstructed RNA molecules are shown from one experiment with 369 mouse fibroblasts, with molecules shown in descending order. **(h)** Sashimi plots visualizing two reconstructed RNA transcripts that supported two distinct transcript isoforms of *Cox7a2l* (ENSMUST00000167741 in orange, and ENSMUST00000025095 in light blue), observed in a mouse fibroblasts (cell barcode: TTCCGTTCGCGACTAA). (**i**) Violin plots showing the percentage of detected molecules that could be assigned to a specific Ensembl transcript isoform, per F1 CAST/EiJ × C57/Bl6J mouse fibroblast. Reported are the results on all Ensembl genes, or the subset with two or more annotated isoforms (‘multi-isoform genes’). The median percentages of assigned molecules per cell were 52.37% and 41.04% for all and multi-isoform genes, respectively. **(j)** Visualizing significant strain-specific isoform expression in mouse fibroblasts, colored by chromosomes. Y-axis shows Benjamini-Hochberg corrected p-values (− log10) from individual Chi-square tests performed per gene evaluating association between allelic origin and isoforms. **(k)** Visualizing the significant strain-specific isoform expression of *Hcfc1r1* in CAST/EiJ and C57/Bl6J mouse strains. Violin plots depict isoform expression in mouse fibroblasts, separated per strain and isoform. Top shows the transcript isoform structures.

We have previously shown that allele-resolved scRNA-seq can be used to infer bursting kinetics of gene expression that are characteristic of transcription^12^. Strikingly, Smart-seq3 based analysis enabled kinetic inference for thousands more genes than using Smart-seq2 alone with a 5’ UMI (11,766 using Smart-seq3; 8,464 using Smart-seq2-UMI) and with significantly improved correlation between the CAST and C57 alleles (0.94 and 0.75 for Smart-seq3 and 0.79 and 0.68 for Smart-seq2-UMI, respectively for burst frequency and size) (**Figure 2f** and **Figure S5**). We conclude that Smart-seq3 enables more sensitive reconstruction of transcriptional bursting kinetics across single cells.

We investigated the lengths of RNAs reconstructed to what extent they contained information on transcript isoform structures. In our experiment with 369 cells, we observed in total 22,196 molecules reconstructed to a length of 1.5kb or longer, and around 200,000 molecules reconstructed to 1kb or longer (**Figure 2g**). Per cell, 8,710 molecules were reconstructed to a length of 500 bp or longer. Importantly, reconstructed molecules could often be assigned to specific transcript isoforms, here exemplified by Sashimi plots for two reconstructed molecules from the *Cox7a2l* gene (**Figure 2h**), which illustrate how reconstructed sequences overlaying exons and splice junctions could assign molecules to transcript isoforms. Intriguingly, 53% of all reconstructed molecules could be assigned to a single annotated Ensembl isoform, including 41% of all molecules detected from multi-isoform genes (**Figure 2i**), thus enabling counting of RNAs at isoform resolution.

Strain-specific transcript isoform regulation has previously been hard to study, since the simultaneously quantification of strain-specific SNPs and splicing outcomes on the same RNAs have not been possible with traditional single-cell or population-level RNA-sequencing. We assigned the *in silico* reconstructed molecules to both allelic origin and transcript isoform structures, which revealed statistically significant strain-specific (CAST or C57) expression of transcript isoforms for 2,172 genes (adjusted p-value < 0.05, chi-square test with Benjamini-Hochberg correction; and p-value < 0.05, gene-specific permutation test) (**Figure 2j** and **Table S2**). For example, transcripts for *Hcfc1r1* were processed into two isoforms (ENSMUST00000024697 and ENSMUST00000179928) that differed both in coding sequence (3 amino acid deletion from a 12-bp alternative 3’ splice site usage) and in 5’ untranslated region splicing. Strikingly, the two isoforms had a significant mutually exclusive pattern of expression between strains (adjusted p-value < 10^−208^, chi-square test with Benjamini-Hochberg correction) (**Figure 2k**). Thus, Smart-seq3 can simultaneous quantify genotypes and splicing outcomes, here exemplified by strain-specific splicing patterns in mouse.

Next, we sought out to benchmark Smart-seq3 on a more complex sample consisting of many different types of cells. To this end, we sequenced 5,376 individual cells from the HCA benchmarking sample^4^, a cryopreserved and complex cell sample comprised of human peripheral blood mononuclear cells (PBMC), primary mouse colon cells and cell line spike-ins of human HEK293T, mouse NIH3T3 and dog MDCK cells. Smart-seq3 cells clearly separated according to species (**Figure S6**) and cell types (**Figure 3a**), and 77% of cells passed quality filtering, significantly higher percentages than the 29% to 63% reported for available protocols^4^, showcasing the robustness of Smart-seq3 (**Figure S7**).

**Figure3.**
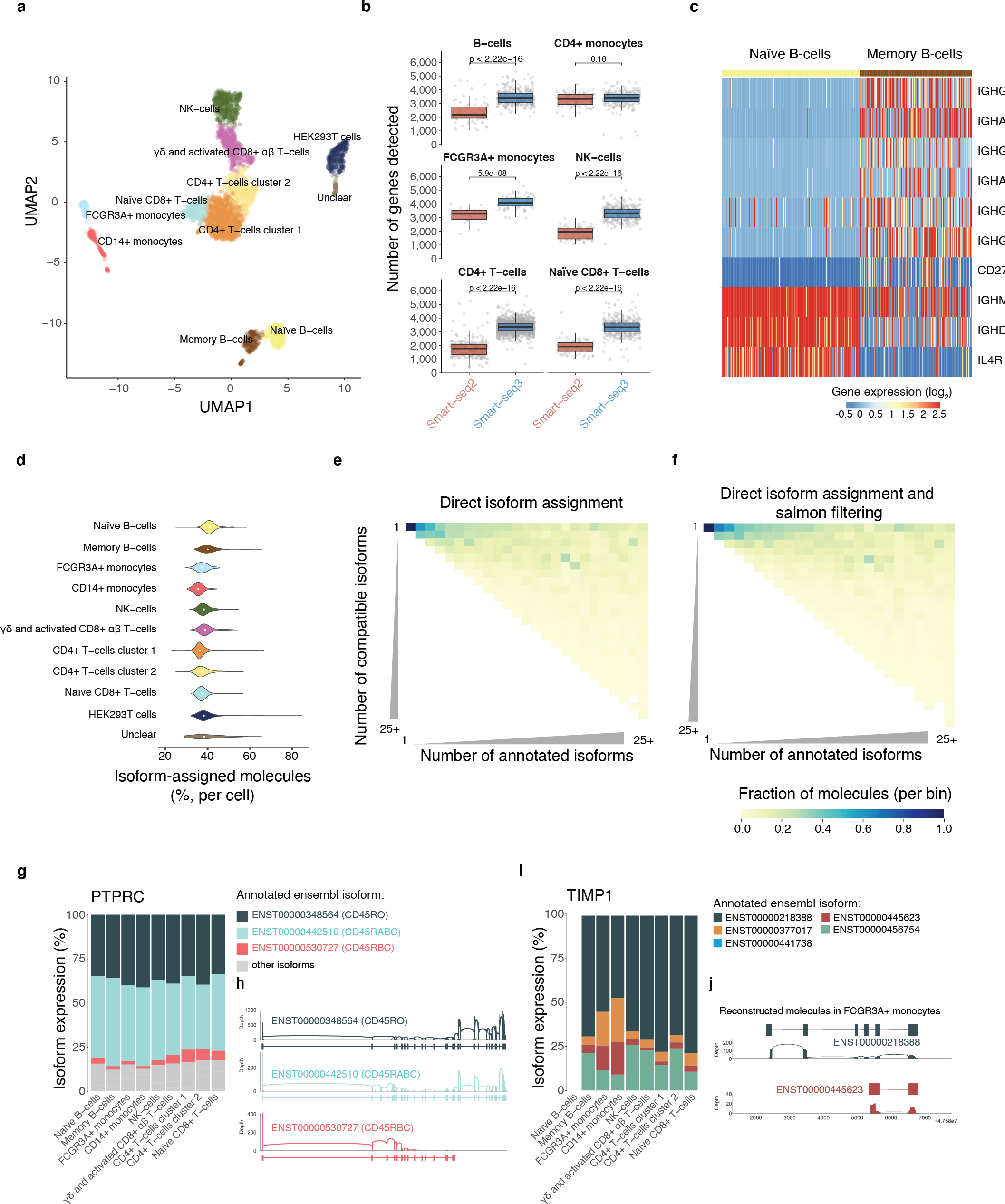
Smart-seq3 analysis of a complex human sample. **(a)** Dimensionality reduction (UMAP) of 3,890 human cells sequenced with the Smart-seq3 protocol and colored by annotated cell type. **(b)** Comparison of sensitivity to detect genes between Smart-seq2 and Smart-seq3 in various cell types. Cells were down-sampled to 100k raw reads per cell and t-test p-values are annotated for each pair-wise comparison. **(c)** Heatmap showing gene expression for selected marker genes that were expressed at statistically significantly different levels in naïve and memory B-cells. Color scale represents normalized and scaled expression values. (**d**) The percentage of reconstructed RNA molecules that could be assigned to a single Ensembl isoform, separated by cell types. (**e**) Matrix showing the fraction of reconstructed molecules that could be assigned to either one or N number of isoforms, where molecules were first grouped by the number of annotated isoform available for its genes. (**f**) Matrix showing the fraction of reconstructed molecules that could be assigned to either one or N number of isoforms (as in e) after we filtered the assignments to only those isoforms with detectable expression (TPM>0) in Salmon (including internal reads without linked UMIs). (**g**) Barplots showing the fraction of molecules assigned to different PTPRC isoforms, separated by cell type and aggregating over all cells within cell types. (**h**) Sashimi plots of reconstructed molecules assigned to either the R0 or RABC isoform of PTPRC in gamma-delta T-cells. (**i**) Barplots showing the fraction of molecules assigned to different TIMP1 isoforms, separating by cell type and aggregating over cells within cell types. (**j**) Sashimi plots of reconstructed molecules assigned to two TIMP1 isoforms in FCGR3A+ monocytes.

Except for CD14+ monocytes, which may be more vulnerable to the year-long freezer storage prior to FACS cell sorting and Smart-seq3 profiling, gene detection sensitivity was significantly higher in all cell types compared to Smart-seq2 already at shallow sequencing depths (**Figure 3b**). This improvement in the number of genes detected extended into traditionally difficult cell types with low mRNA content, such as T-cells and B-cells for which we typically observed one thousand more genes per cell. Interestingly, we detected two distinct clusters of B-cells (**Figure 3a**) that were not separated in single-cell data from existing methods^4^. Differential expression between the B-cell populations reported 279 genes with significant expression difference, which included several known marker genes for naïve and memory B cells (**Figure 3c**). This demonstrated an improved ability of Smart-seq3 to separate biologically meaningful clusters of cells compared to existing methods.

Investigating the RNA molecule reconstruction performance across the human cell types, revealed that 36-41% of all detected molecules could be assigned to a specific isoform across cell types (**Figure 3d**). To investigate the isoform assignment in greater detail, we visualized the number of compatible isoforms for each reconstructed RNA molecule, binning genes by the number of annotated isoforms. Many additional molecules could be assigned to a small set of transcript isoforms (**Figure 3d**). We further reasoned that the internal reads in Smart-seq3 could provide more information on isoform expression. To this end, we computed isoform expressions using Salmon^15^ on all reads from Smart-seq3 and filtered the direct RNA reconstruction based assignment of molecules to only those isoforms that had detectable expression (TPM>0) in Salmon (**Methods**). This strategy further increased the assignment of molecules to unique isoforms (42% of all molecules) (**Figure 3e**), and we used the Salmon-filtered isoform expression levels for the remainder of the study.

Next, we investigated the patterns of isoform expression across cell types. Strikingly, 2,186 genes had statistically significant patterns of isoform expressions across cell-types (Adjusted p-values <0.05; Kruskal-Wallis test and Benjamini-Hochberg correction). (**Table S3**). One of the significant genes was PTPRC (also known as CD45) which can be post-transcriptionally processed into several different isoforms^16^, including a full-length isoform (called RABC) and one that has excluded three consecutive exons (called RO). We mainly observed these two isoforms across the human immune cell types, although at significantly varying levels (**Figure 3f**). Aggregating the reads supporting these two isoforms in gamma-delta T-cells (**Figure 3g**) further shows how the reconstructed molecules separated the inclusion or skipping of the three consecutive exons. Other specific isoform patterns were shared by certain cell types, for example both CD14+ and FCGR3A+ monocytes expressed specific isoforms of the TIMP1 gene (**Figure 3h,i**). Both monocyte populations specifically expressed a shorter isoform of the TIMP1 gene, whereas the long, full-length isoform was dominant across other cell types (**Figure 3h**), again supported by the reconstructed molecules (**Figure 3i**). Altogether, these results highlight the new and unique capabilities of using Smart-seq3 to query isoform expression and regulation across cell types.

## Discussion

Mammalian genes typically produce multiple transcript isoforms from each gene^17^, with frequent consequences on RNA and protein functions. Analysis of transcript isoform expression (in single cells or in cell populations) using short-read sequencing technologies have often focused on individual splicing events (e.g. skipped exon) or used the read coverage over shared and unique isoform regions to infer the most likely isoform expression^18,19^. This is due to paired short reads seldom having sufficient information to assess interactions between distal splicing outcomes or combined with allelic expression from transcribed genetic variation. Long-read sequencing technologies can used to directly sequence transcript isoforms in single cells^2,3^. However, these strategies have limited cellular throughput and depth. For example, the Mandalorion approach provided comprehensive isoform data for seven cells^2^, whereas scISOr-seq investigated isoform expression in thousands of cells at an average depth of 260 molecules per cell^3^. In contrast, we obtained on average 8,710 reconstructed molecules per cell (above 500 bp). Moreover, in scISOr-seq the pre-amplified cDNA was sequenced on both short- and long-read sequencers in parallel to characterize cell types and sub-types, and the isoform-level sequencing data was mainly aggregated over cells according to clusters^3^. The use of two parallel library construction methods and sequencing technologies for the same pre-amplified cDNA from individual cells substantially increases cost and labor.

We developed Smart-seq3 to be both highly sensitive, thus improving the ability to identify cell types and states, and isoform-specific, to simultaneously reconstruct millions of partial transcripts across cells. Smart-seq3 thus removes the additional costs and labor associated with the use of multiple library preparation technologies and sequencing platforms in parallel. Compared to known transcript isoform annotations, these partial transcript reconstructions were sufficient to assign 40-50% of detected molecules to a specific isoform, which further revealed strain- and cell-type specific isoform regulation. Excitingly, this reconstruction should improve the abilities to perform splicing quantitative trait loci mapping, since both splicing outcomes and transcribed SNPs can now be directly quantified. The full Smart-seq3 protocol has been deposited at protocols.io (dx.doi.org/10.17504/protocols.io.7dnhi5e) and can be readily implemented by molecular biology laboratories without the need for specialized equipment.

Several large-scale projects aim to systematically construct cell atlases across human tissues and those of model organisms^20^. These efforts are increasingly relying on scRNA-seq methods that count RNAs towards annotated gene ends (e.g. 10× genomics) that provides little information on isoforms expression patterns across cell types and tissues. Moreover, large-scale efforts are also emerging to use single-cell genomics for the systematic analysis of disease (e.g. the LifeTime project) to identify disease mechanisms and consequences. As post-transcriptional gene regulation has been tightly linked to disease^21^, it would be a missed opportunity for such efforts and atlases to disregard isoform-level expression patterns. In contrast to long-read sequencing efforts, Smart-seq3 simultaneously provides cost effective gene expression profiling across cell types and isoform-resolution RNA counting within the same assay. This is currently achieved at a cost per sequence ready cell library around 0.5-1 EUR. Additionally, as the current implementation uses 384-well plates, it is also possible to first shallowly sequence all cells and then later select cells of rare cell populations (as cellular amplified cDNAs can be kept in individual wells for extended periods of time) for in-depth sequencing and transcript isoform reconstruction. Altogether, we introduced a scRNA-seq method that is applicable to characterize cell types and annotate cell atlases at the level of gene, isoform and allelic expression.

## Acknowledgements

We would like to thank Holger Heyn for providing us with the HCA sample, Swati Parekh for help with the zUMIs pipeline and Björn Reinius for discussions. C.Z. is supported by an EMBO long-term fellowship (ALTF 673-2017). G-J.H. is funded by HFSP long-term fellowship LT000155/2017-L. This work was supported by grants to R.S. from the European Research Council (648842), the Swedish Research Council (2017-01062), the Knut and Alice Wallenberg’s foundation (2017.0110), the Bert L. and N. Kuggie Vallee Foundation, the Göran Gustafsson Foundation, the National Institute of Health.

## Competing financial interests

R.S., M.H-J. and O.R.F have filed a patent application on Smart-seq3.

## Data and code availability

All sequencing data will be deposited in the European Nucleotide Archive (ENA) at the European Bioinformatics Institute (EBI). Processing of Smart-seq3 libraries has been incorporated in zUMIs (https://github.com/sdparekh/zUMIs) and additional code for molecule reconstruction will be made available through github (https://github.com/sandberg-lab).

## Author contribution

Developed Smart-seq3 chemistry: M.H-J and O.R.F, with input from C.Z. Generated single-cell RNA-seq libraries: M.H-J. Developed the reconstruction procedure: C.Z., P.C., with input from M.H-J. and G-J.H. Performed computational analysis: C.Z., P.C., M.H-J., with input from D.R. and A.L. Prepared figures: M.H-J., C.Z., P.C. and R.S. Planned and supervised the overall work: R.S. Wrote the manuscript: R.S, with input from C.Z. and M. H-J.

## Supplementary Figures

**Supplementary Figure 1.**
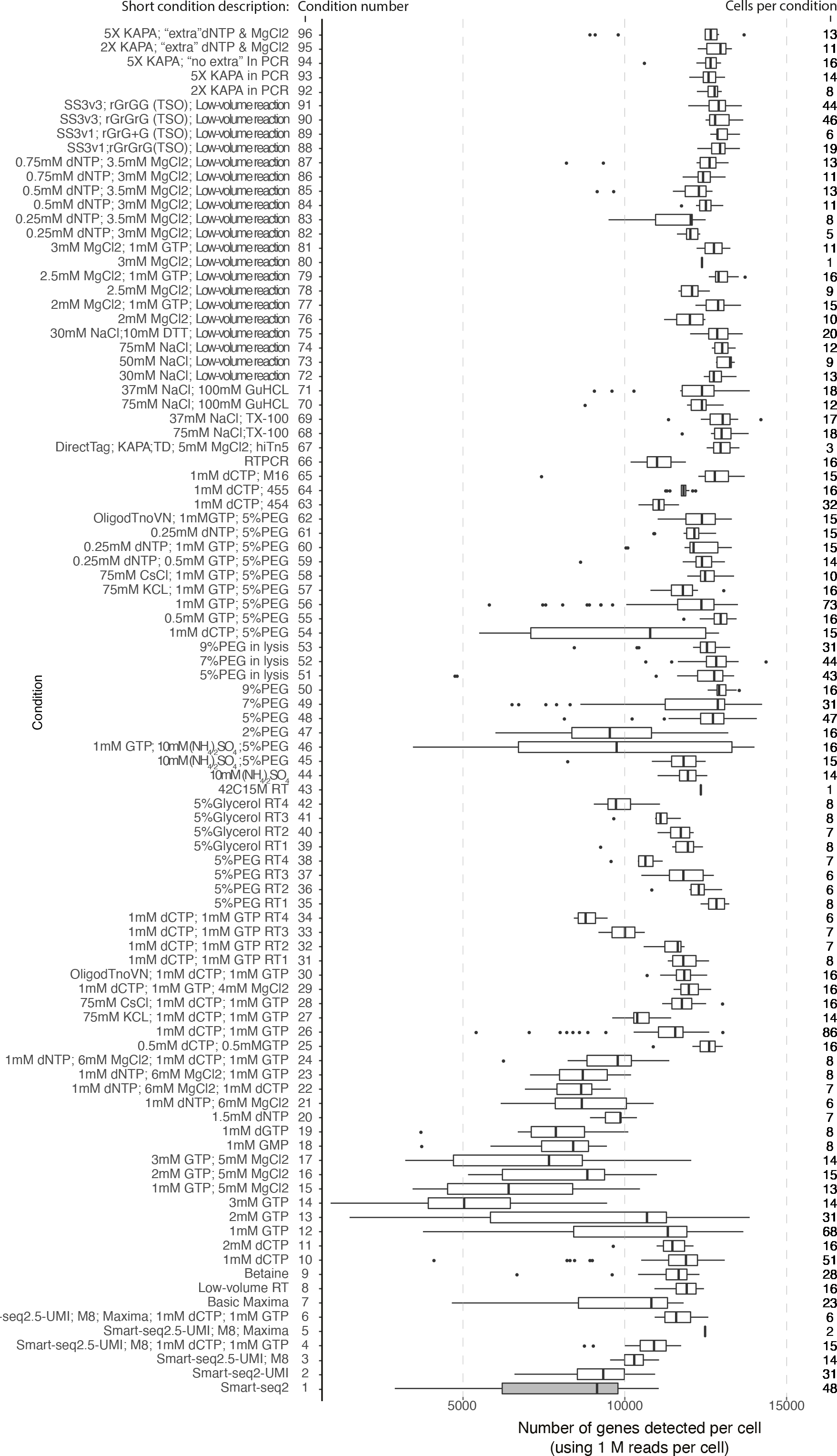
Overview of sequenced conditions and iterations of Smart-seq3. Each row shows a tested reaction condition and the number of genes detected in individual HEK293FT cells at 1M raw fastq reads. The numbers of individual cells that contained at least one million sequenced reads per condition are listed on the right. Several earlier versions of Smart-seq2 with elements of Smart-seq3 chemistry are included as “Smart-seq2.5” in this figure. The exact reaction conditions per row are listed in Supplementary Table 1.

**Supplementary Figure 2.**
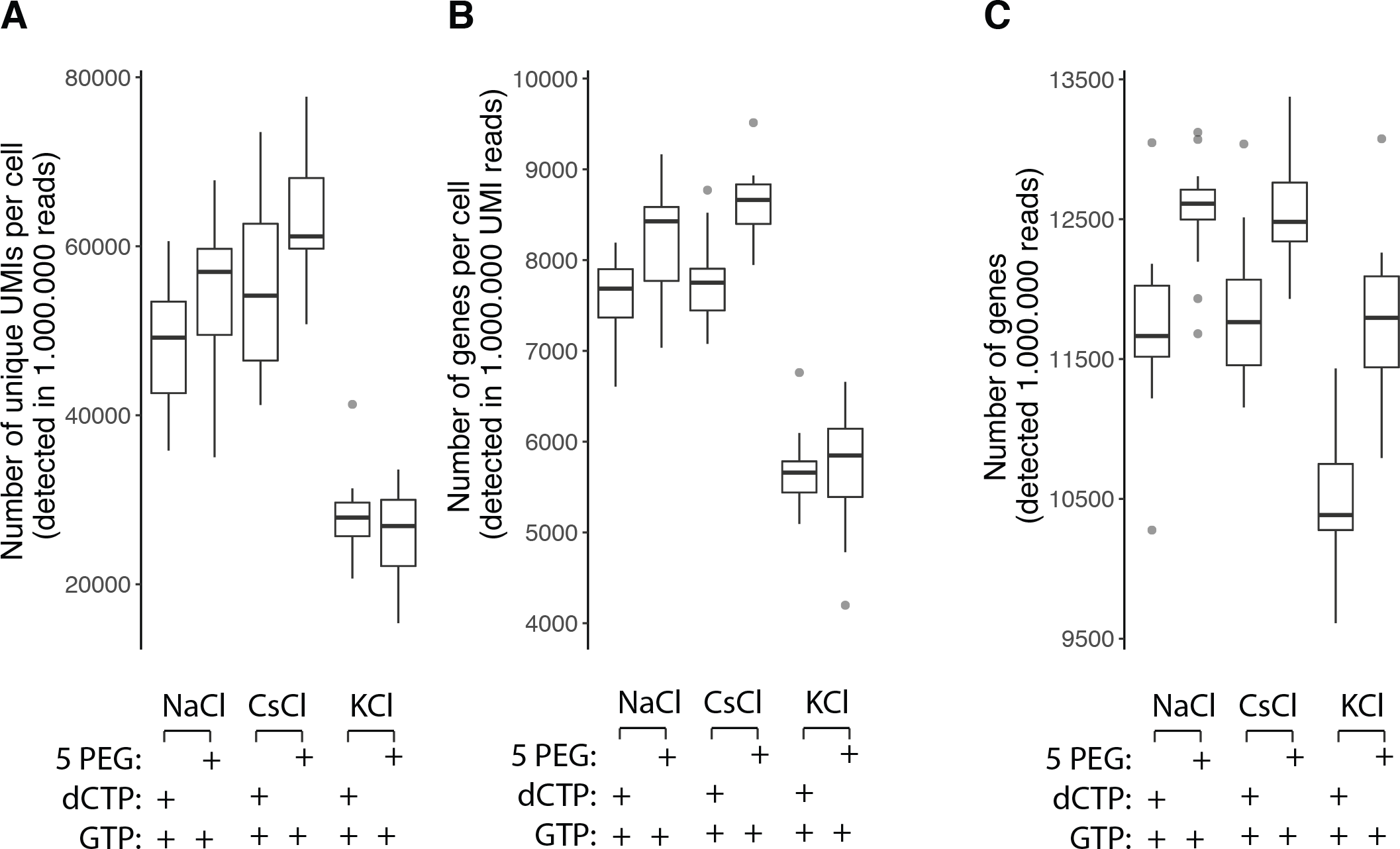
Effects of salts, PEG and additives on Smart-seq3 reverse transcription. (**a**) Testing the performance of Maxima H-minus reverse transcription reactions on different reaction conditions. For each condition, we summarized boxplots with the number of unique UMIs detected in individual HEK293FT cells at 1M raw fastq reads. We tested reverse transcription in the context of using a NaCl, CsCl or the standard KCl based buffer. Moreover, we evaluated the effects of adding of 5% PEG or 1mM dCTP (16 cells per condition). (**b**) Reaction conditions as in (a) summarized against the number of genes identified from 1 million raw UMI-reads per cell (16 cells per condition). (**c**) Reaction conditions as in (a) summarized against the number of genes identified from 1 million raw reads (sub-sampling from both 5’ UMI and internal reads) per cell (16 cells per condition).

**Supplementary Figure 3:**
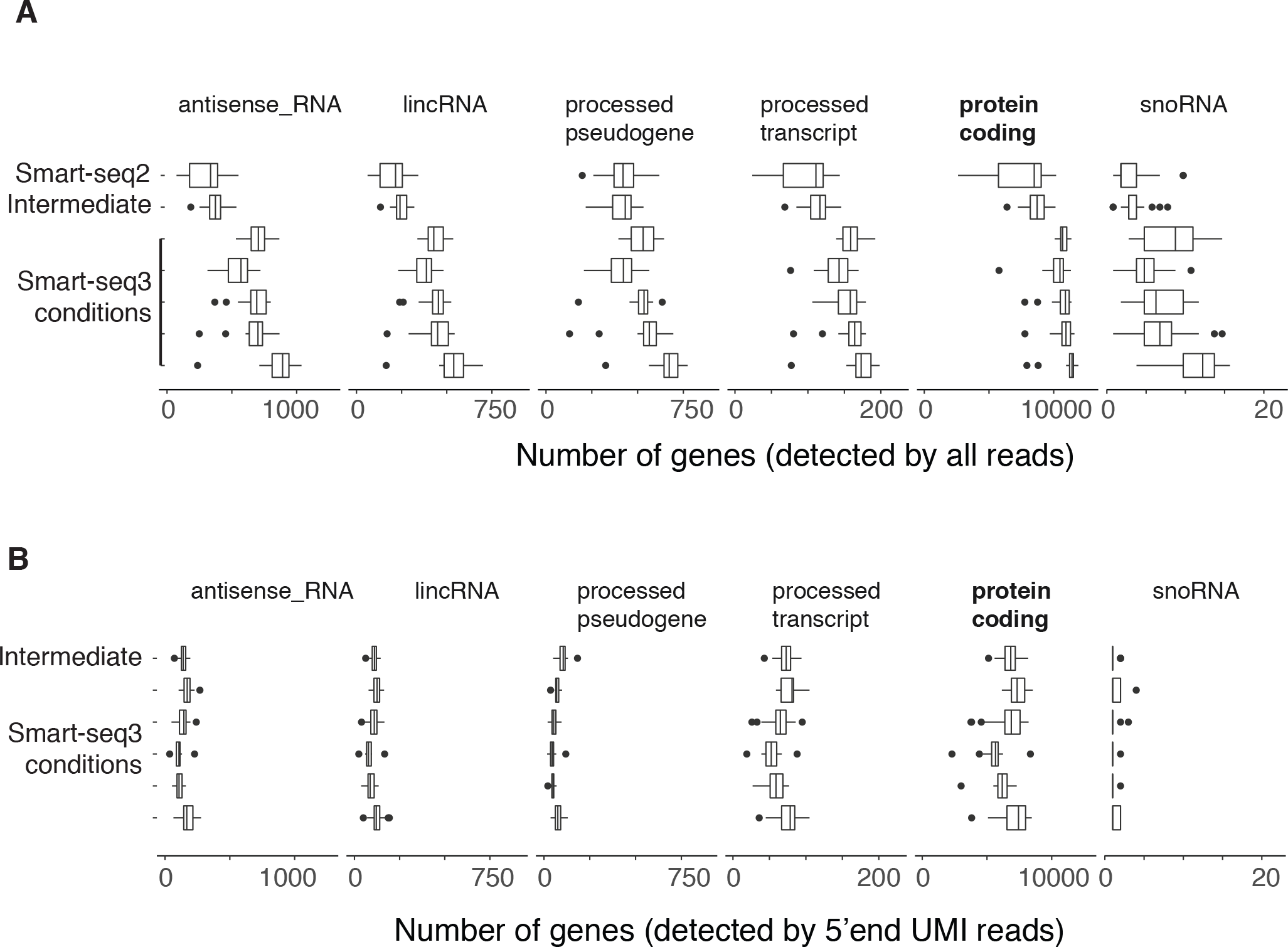
Improved detection of protein-coding and non-coding RNAs with Smart-seq3. (**a**) Variants of Smart-seq3 reactions show improved detection of protein coding genes and also genes of different biotypes, including poly-A+ lincRNAs, antisense RNAs, processed pseudogenes, processed transcripts and snoRNAs, compared to Smart-seq2 and earlier experimentations of Smart-seq2 with UMIs (here called “intermediate”). (**b**) Shows genes detected of similar RNA biotypes by UMI containing reads in Smart-seq2 with UMIs (here called “intermediate”) and Smart-seq3 variants.

**Supplementary Figure 4:**
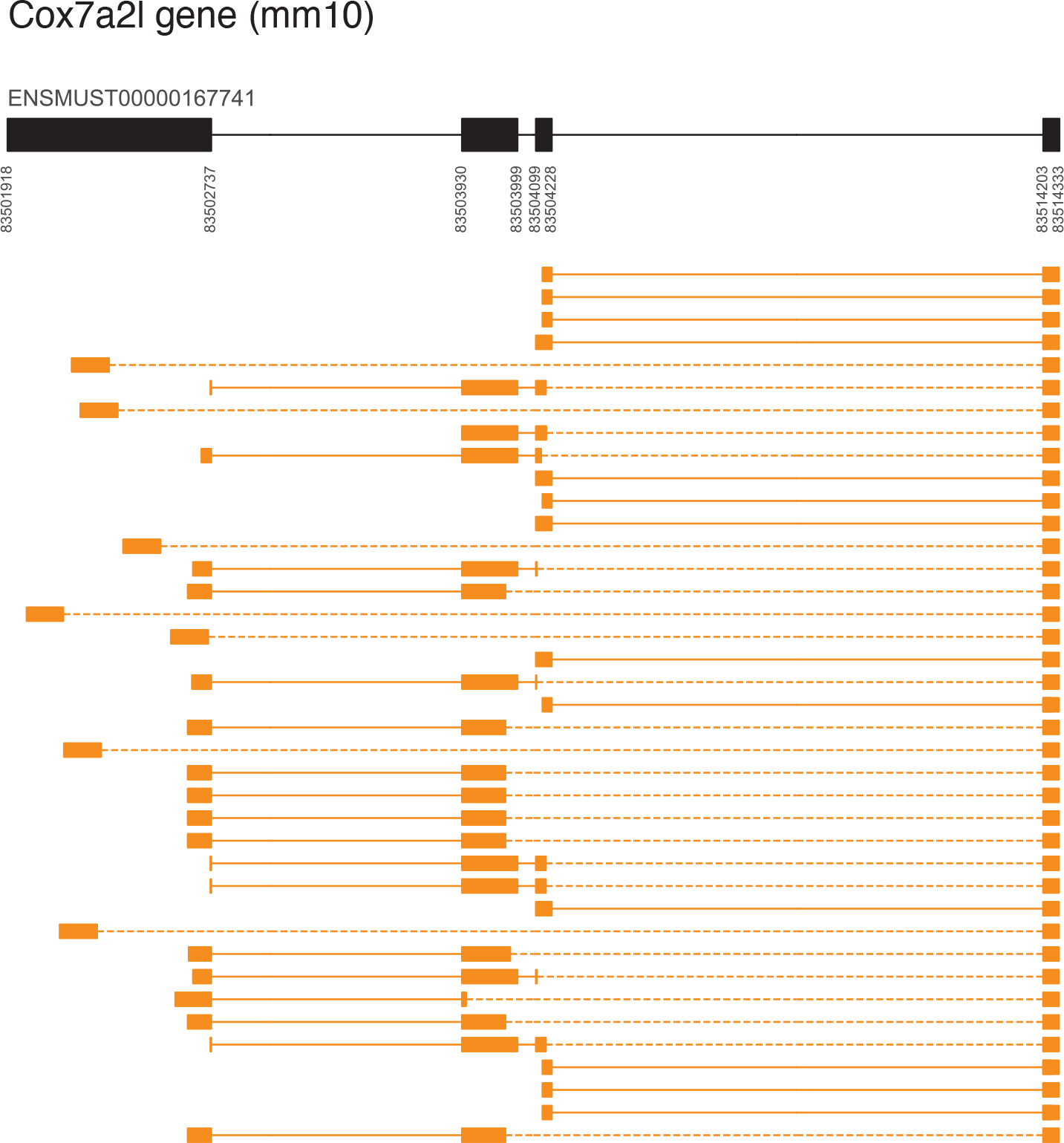
Visualization of read-pairs from a single transcribed molecule from Cox7a2 locus in primary fibroblast cell. Visualization of read pairs sequenced from one molecule from the Cox7a2l locus. Top show the exons and introns in the Cox7a2l locus, with genomic coordinates (mm10). Each row show a unique read pair, where oranges boxes show the mapping of sequences onto the genomic loci, dotted lines indicate that the sequences are connected by the read pairs and solid lines represent that the exon-intron junction was captured in the sequenced reads. Note, all read pairs combined span essentially the full transcript, meaning that for this molecule we could reconstruct the full transcript.

**Supplementary Figure 5:**
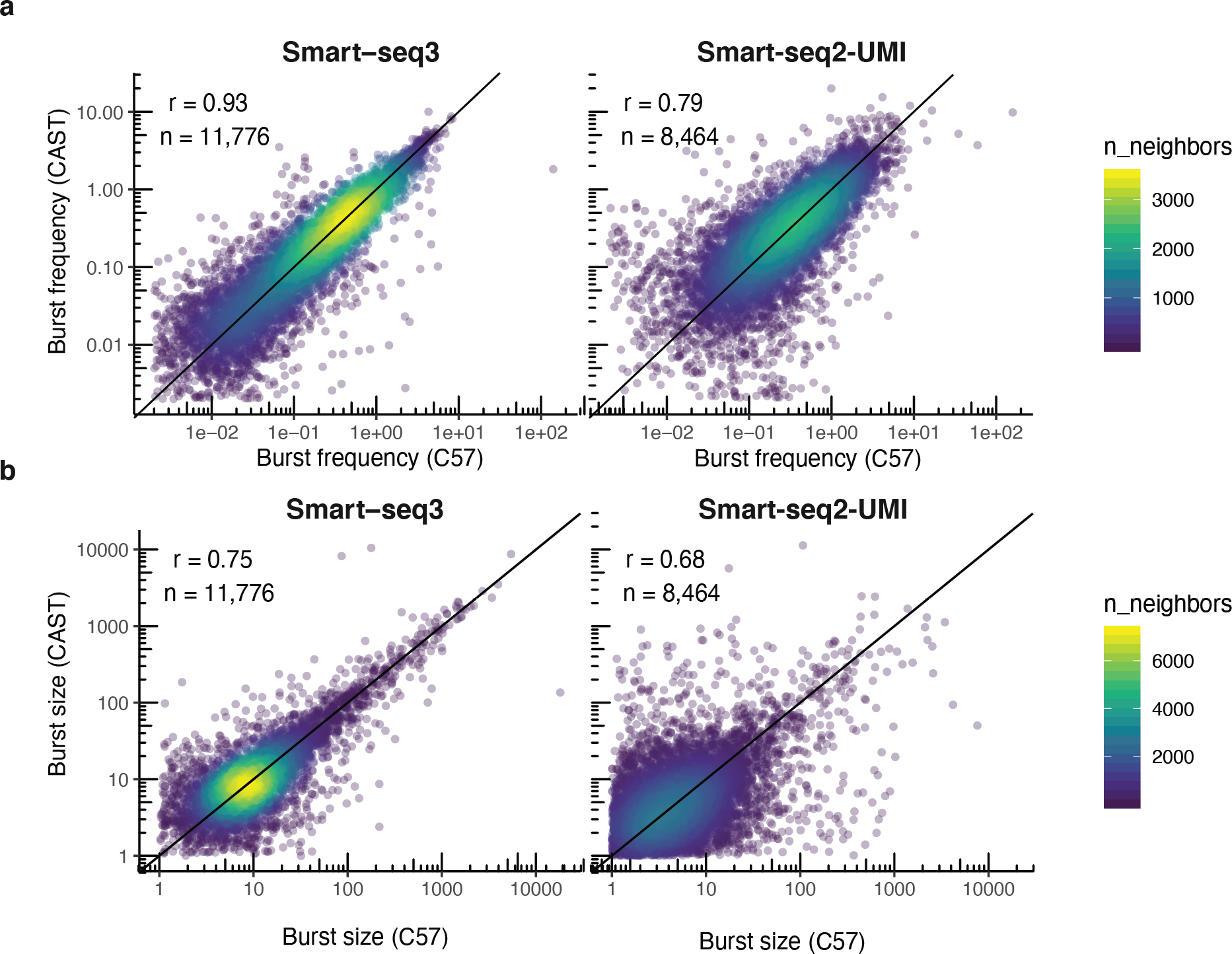
Detailed comparison of burst kinetics inference based on Smart-seq2-UMI and Smart-seq3 data. (**a**) Scatter plots showing the burst frequencies inferred for the C57 (x-axis) and CAST (y-axis) alleles for genes in mouse fibroblasts. The left plot show the results based on Smart-seq3 data and the right panel show the results from using Smart-seq2-UMI data. (**b**) Scatter plots showing the burst sizes inferred for the C57 (x-axis) and CAST (y-axis) alleles for genes in mouse fibroblasts. The left plot show the results based on Smart-seq3 data and the right panel show the results from using Smart-seq2-UMI data.

**Supplementary figure 6:**
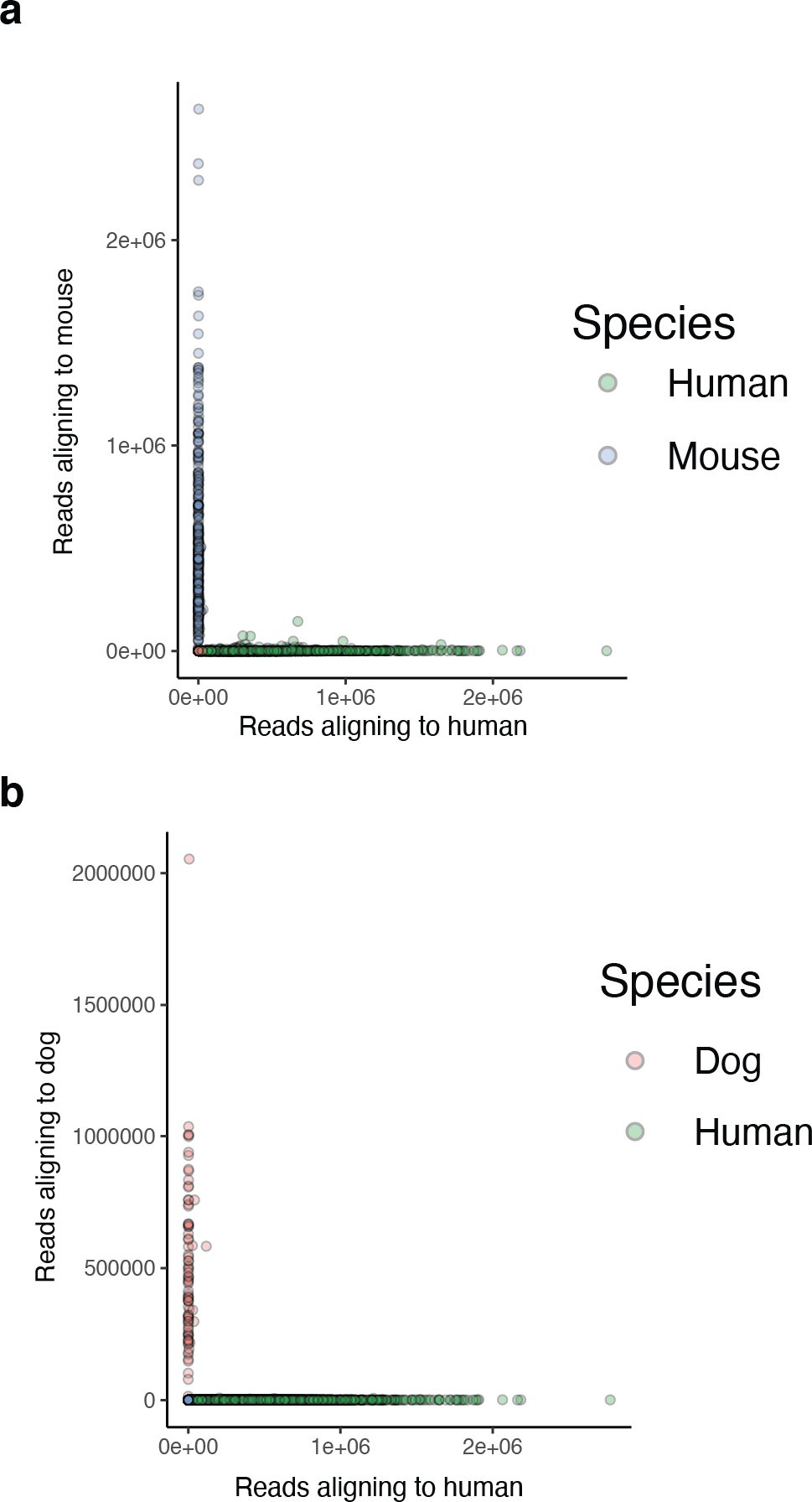
Species-mixing and doublets in Smart-seq3. (**a**) Scatter plot showing the number of reads that aligned to human (x-axis) and mouse (y-axis) for the complex HCA sample that contained both human, mouse and dog cells. (**b**) Scatter plot showing the number of reads that aligned to human (x-axis) and dog (y-axis) for the complex HCA sample that contained both human, mouse and dog cells. Few cells show any signal towards more than on genome, demonstrating a very low doublet rate.

**Supplementary figure 7:**
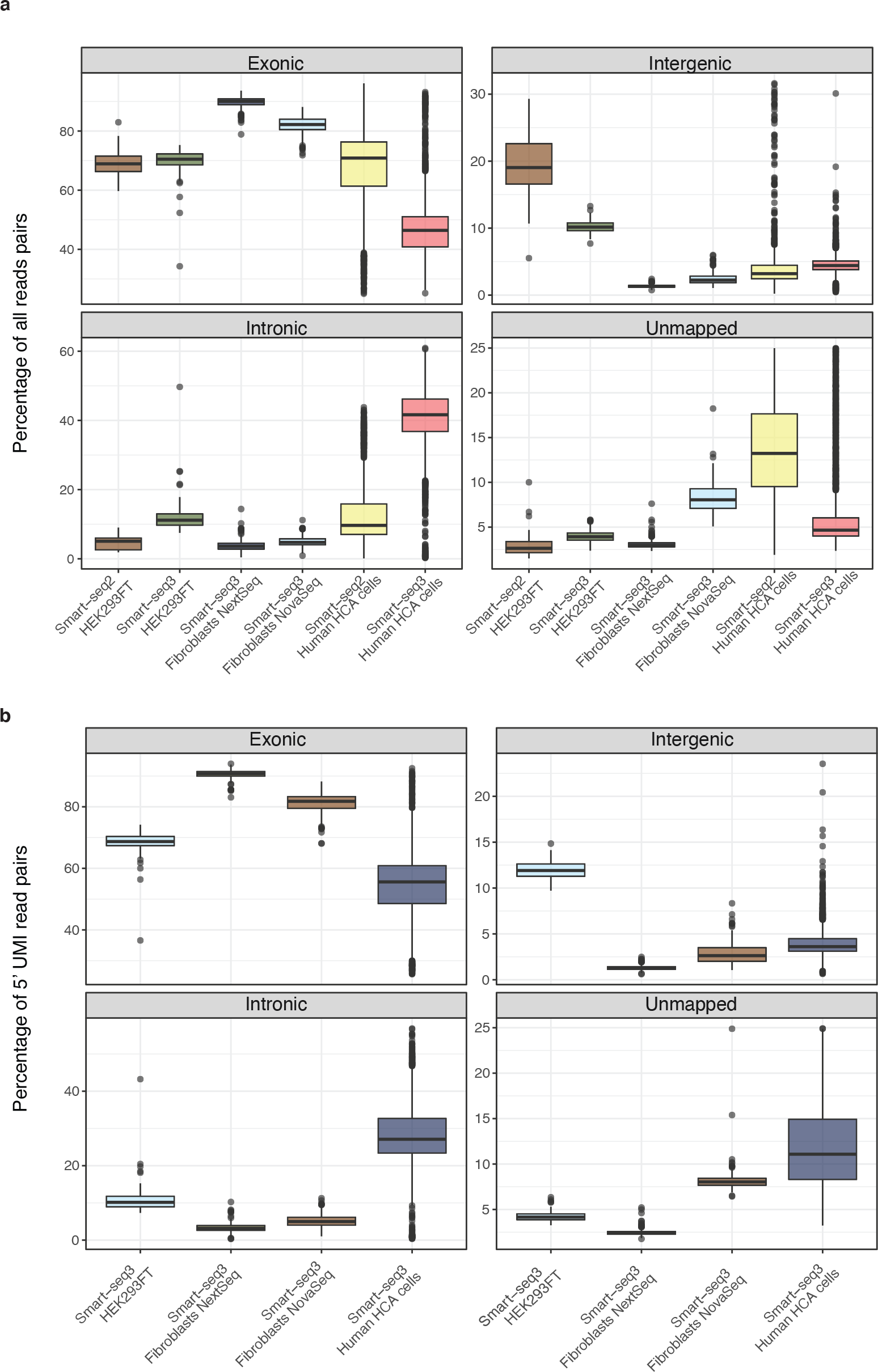
Mapping statistics of used Smart-seq2 and Smart-seq3 libraries. (**a**) Percentage of unmapped read pairs, and read pairs that aligned to exonic, intronic and intergenic regions. Separated per protocol (Smart-seq2 and Smart-seq3) and experiment (HEK293FT, Mouse Fibroblasts, HCA cells). (**b**) Mapping statistics for 5’UMI-containing read pairs in Smart-seq3. Percentage of unmapped read pairs, and read pairs that aligned to exonic, intronic and intergenic regions. Separated per experiment (HEK293FT, Mouse Fibroblasts, HCA cells).

## Supplementary Methods

### Cell cultures

HEK293FT cells (Invitrogen) were cultured in complete DMEM medium containing 4.5g/L glucose and 6mM L-glutamine (Gibco), supplemented with 10% Fetal Bovine Serum (Sigma-Aldrich), 0.1mM MEM Non-essential Amino Acids (Gibco), 1mM Sodium Pyruvate (Gibco) and 100 μg/mL Pencillin/Streptomycin (Gibco). Cells were dissociated using TrypLE express (Gibco) and stained with Propidium Iodide, to exclude dead cells, before distribution into 96 or 384 well plates containing 3μL lysis buffer using a BD FACSMelody 100 μm nozzle (BD Bioscience). The Smart-seq3 lysis buffer consisted of 0.5 unit/μL Recombinant RNase Inhibitor (RRI) (Takara), 0.15% Triton X-100 (Sigma), 0.5mM dNTP/each (Thermo Scientific), 1μM Smart-seq3 oligo-dT primer (5’-Biotin-ACGAGCATCAGCAGCATACGA T_30_VN-3’; IDT), 5% PEG (Sigma) and 0.05 μL of 1:40.000 diluted ERCC spike-in mix 1 (For HEK293FT cells). The plates were spun down immediately after sorting and stored at −80 degrees.

Primary mouse fibroblasts were obtained from tail explants of CAST/EiJ × C57/Bl6J derived adult mice (with ethical approval from the Swedish Board of Agriculture, Jordbruksverket: N343/12). Cells were cultured and passaged twice in (DMEM high glucose (Invitrogen), 10% ES cell FBS (Gibco), 1% Penicillin/Streptomycin (Invitrogen), 1% Non-essential amino acids (Invitrogen), 1% Sodium-Pyruvate (Invitrogen), 0.1mM b-Mercaptoethanol (Sigma), before stained with Propidium Iodide, and sorted in to 384 well plates containing 3μL Smart-seq3 lysis buffer. Again, plates were spun down and stored at −80 degrees immediately after sorting.

The Human Cell Atlas (HCA) reference sample consisting of a mix of Human PBMCs, Mouse colon, as well as fluorescent labelled cell-lines HEK-293-RFP, NiH3T3-GFP and MDCK-Turbo650 were thawed according to specified instructions^4^. Cells were stained with Live/Dead fixable Green Dead cell stain kit (Invitrogen), facilitating the exclusion of dead cells as well as NIH3T3-GFP cells. Additionally, both debris and doublets were excluded in the gating. Cells were index sorted into 384 well plates, containing 3μL Smart-seq3 lysis buffer, using a BD FACSMelody sorter with 100μm nozzle (BD Bioscience).

### Generation of Smart-seq2 libraries

Smart-seq2 cDNA libraries were generated according the published protocol^22^. For Smart-seq2-UMI, cDNA libraries were generated as previously published^12^. Recipes for other “intermediate” Smart-seq2 reactions can be found in Table S1. Tagmentation was performed with similar cDNA input and volumes as for Smart-seq3 described below.

### Generation of Smart-seq3 libraries

To facilitate cell lysis and denaturation of the RNA, plates were incubated at 72 degrees for 10 min, and immediately placed on ice afterwards. Next, 1μL of reverse transcription mix, containing 25 mM Tris-HCL pH 8.3 (Sigma), 30 mM NaCl (Ambion), 1 mM GTP (Thermo Scientific), 2.5 mM MgCl2 (Ambion), 8 mM DTT (Thermo Scientific), 0.5 u/μL RRI (Takara), 2 μM of different Smart-seq3 Template switching oligo (TSO) (see additional table for list of evaluated TSOs; 5’-Biotin-AGAGACAGATTGCGCAATGNNNNNNNNrGrGrG-3’; IDT) and 2 u/μL Maxima H-minus reverse transcriptase enzyme (Thermo Scientific), were added to each sample. Reverse transcription and template switching were carried out at 42 degrees for 90min followed by 10 cycles of 50 degrees for 2min and 42 degrees for 2 min. The reaction was terminated by incubating at 85 degrees for 5 min. PCR preamplification was performed directly after reverse transcription by adding 6 μL of PCR mix, bringing reaction concentrations to 1x KAPA HiFi PCR buffer (contains 2mM MgCl2 at 1×) (Roche), 0.02u/μl DNA polymerase (Roche), 0.3mM dNTPs, 0.1μM Smartseq3 Forward PCR primer (5’-TCGTCGGCAGCGTCAGATGTGTATAAGAGACAGATTGCGCAATG-3’; IDT), 0.1μM Smartseq3 Reverse PCR primer (5’-ACGAGCATCAGCAGCATACGA-3’; IDT). PCR was cycled as follows: 3min at 98 degrees for initial denaturation, 20-24 cycles of 20 secs at 98 degrees, 30 sec at 65 degrees, 6 min at 72 degrees. Final elongation was performed for 5 min at 72 degrees. For various iterations and optimization conditions, see Supplementary table 1 for information about specific conditional changes to library preparation.

### Sequence library preparation

Following PCR preamplification, all samples, regardless of protocol used, were purified with either AMpure XP beads (Beckman Coulter) or home-made 22% PEG beads (see step 27 in protocol doi:10.17504/protocols.io.p9kdr4w at protocols.io). Library size distributions were checked on a High sensitivity DNA chip (Agilent Bioanalyzer) and all cDNA concentrations were quantified using the Quant-iT PicoGreen dsDNA Assay Kit (Thermo Scientific). cDNA was subsequently diluted to 100-200pg/uL. Tagmentation was carried out in 2 uL, consisting of 1× tagmentation buffer (10mM Tris pH 7.5, 5mM MgCl2, 5% DMF), 0.08-0.1 uL ATM (Illumina XT DNA sample preparation kit) or TDE1 (Illumina DNA sample preparation kit), 1 uL cDNA and H2O. Plates were incubated at 55 degrees for 10min, followed by addition of 0.5 uL 0.2% SDS to release Tn5 from the DNA. Library amplification of the tagmented samples was performed using either 1.5 uL Nextera XT index primers (Illumina) or 1.5 uL custom designed Nextera index primers containing either 8 or 10 bp indexes (0.1 uM each), differing with a minimal levenshtein distance of 2 between any two indices. 3 uL PCR mix (1× Phusion Buffer (Thermo Scientific), 0.01 U/uL Phusion DNA polymerase (Thermo Scientific), 0.2 mM dNTP/each) was added to each well, and incubated at 3 min 72 degrees; 30 sec 95 degrees; 12 cycles of (10 sec 95 degrees; 30 sec 55 degrees; 30 sec 72 degrees); 5 min 72 degrees; in a thermal cycler. For the experiments optimizing the UMI fragment conditions, following changes to the tagmentation procedure (cDNA input, amount of ATM, and time at 55 degrees) are shown in **Figure 1c**. After tagmentation samples were pooled, and the pool purified with Ampure XP beads or 22% home-made PEG beads at 1:0.6 ratio. Libraries were sequenced at 75 bp single-end, or 150 bp paired-end on a high output flow cell using the Illumina NextSeq500 instrument, or on a NovaSeq S4 flow cell 150 bp paired-end.

### Gel cutting pilot

We additionally experimented with selecting for certain lengths of libraries prior to sequencing of the mouse fibroblast cells. We used 20uL of purified sequence ready library and loaded it onto a 2% Agarose E-Gel EX and ran the gel for 12min. We manually cut the gel in the regions corresponding to 550-2000bp and re-purified the library using Qiagen QiaQuick gel extraction kit following the manufacturers protocol. We observed a modest improvement, however selecting for longer fragments could likely improve reconstruction lengths.

### Read alignments and gene-expression estimation

Raw non-demultiplexed fastq files were processed using zUMIs (version 2.4.1 or newer) with STAR (v2.5.4b), to generate expression profiles for both the 5’ ends containing UMIs as well as combined full length and UMI data. To extract and identify the UMI-containing reads in zUMIs, find_pattern: ATTGCGCAATG was specified for file1 as well as base_definition: cDNA(23-75; Single-end), (23-150bp, paired-end) and UMI(12-19) in the YAML file. UMIs were collapsed using a Hamming distance of 1. Human cells were mapped to hg38 genome and mouse fibroblast cells were mapped against mm10 genome with CAST SNPs masked with N to avoid mapping bias, both supplemented with additional STAR parameters “--limitSjdbInsertNsj 2000000 --outFilterIntronMotifs -- RemoveNoncanonicalUnannotated --clip3pAdapterSeq CTGTCTCTTATACACATCT”. Experiments containing HEK293FT cells were quantified with gene annotations from Ensembl GRCh38.91. Mouse primary fibroblast data was quantified with gene annotations from Ensembl GRCm38.91.

### Allele-calling of F1 mouse molecules

CAST/EiJ strain specific SNPs were obtained from the mouse genome project^23^ dbSNP 142 and filtered for variants clearly observed in existing CAST/EiJ × C57/Bl6J F1 data, yielding 1,882,860 high-quality SNP positions. Uniquely mapped read pairs were extracted and CIGAR values parsed using the GenomicAlignments package^24^. Reads with coverage over known high-quality SNPs were retained and grouped by UMI sequence. Molecules with >33% of bases at SNP positions showing neither the CAST nor the C57 allele were discarded and we required >66% of observed SNP bases within molecules to show one of the two alleles to make an assignment.

### Inference of transcriptional burst kinetics

Allele-resolved UMI counts were used to generate maximum likelihood inference of bursting kinetics from scRNA-seq data as described previously^12^. Inference scripts are available at https://github.com/sandberg-lab/txburst. To ensure a fair comparison with the data generated in this study, we reprocessed the Smart-seq2 data deposited at the European Nucleotide Archive accession E-MTAB-7098 using zUMIs and the same SNP set as described above.

### Primary data processing for mixed-species benchmarking sample

The complete dataset was mapped against a combined reference genome for human (hg38), mouse (mm10) and dog (CanFam3.1). Cells mapping clearly (> 75% of reads) to the mouse or dog were removed. Remaining cells representing HEK293, PBMCs and potential low quality libraries were processed using zUMIs (version 2.5.5) and mapped against the human genome only.

### Analysis of human HCA benchmark samples

First, cells were filtered for low quality libraries requiring >10,000 raw reads, >75% of reads mapped to the genome and >25% exonic fractions. Further analysis was done within v3.1 of Seurat^25^ retaining cell with > 500 genes detected (intron+exon quantification). Data was normalized (“LogNormalize”) and scaled to 10,000 as well as regressing out the total number of counts per cell. The top 2,000 variable genes were found using the “vst” method and used for PCA dimensionality reduction. The first 20 principal components were used for both SNN neighborhood construction as well as UMAP dimensionality reduction. Lastly, louvain clustering was applied (resolution = 0.7) to find cell groupings. Major cell types were readily identifiable by common marker genes: CD4+ T-cells (CD4, IL7R, CD3D, CD3E, CD3G), CD8+ T-cells (CD8A, CD8B), CD14+ Monocytes (CD4, CD14, S100A12), FCGR3A+ Monocytes (FCGR3A), B-cells (MS4A1, CD19, CD79A), NK-cells (NKG7, LYZ, NCAM1) and HEK cells (high number of genes detected). Naïve T-cells were separated from activated by CCR7, SELL, CD27, IL7R and lack of FAS, TIGIT, CD69. γδ T-cells were separated from other T-cells by TRGC1, TRGC2, TRDC and lack of TRAC, TRBC1, TRBC2.

### Isoform reconstruction of UMI-linking fragments from Smart-seq3

The genomic alignments of 5’ UMI containing reads and their paired reads from same fragments were generated by zUMI (version 2.4.1 or newer) with UMI and cell barcode error correction. Unique and multi-mapped reads from same molecules mapping to exonic regions were used for isoform reconstruction. The genomic positions of exons from each isoform were based on reference gene annotation from Ensembl GRCm38.91 for mouse fibroblast data and Ensembl GRCh38.95 for human HCA data. Reads mapping to same molecule were compared to annotated transcripts structures, and represented as a Boolean string indicating which exon were found in read pairs and junctions (“1”) and junctions supporting the exclusion of exons (“0”). For exons not covered with reads, “N” was used to signify lacking. The Boolean string from the reconstructed molecule were matched to the string corresponding to each reference isoforms of same gene to return compatible isoform(s) for each molecule. Molecule isoform assignments were further corrected based on reads aligning to alternative 5’ and 3’ splice sites of overlapping exons from different isoforms.

### Isoform assignments by integrating non-UMI reads

Transcriptome bam files generated using zUMI were demultiplexed per cell and isoform abundances quantified using Salmon^15^ (v0.14.0) quant command and using he following settings “--fldMean 700 --fldSD 100 --fldMax 2000 --minAssignedFrags 1 --dumpEqWeights”. We corrected the Salmon output for cases where all reads were assigned to one out of many possible isoforms belonging to the same equivalent classes. For each cell, isoforms with TPM > 0 from salmon were considered expressed, and used to filter compatible isoforms of the reconstructed molecules.

If more than one isoform was compatible with a reconstructed molecule (after Salmon filtering), each compatible isoform obtained a partial molecule count (1/N compatible isoforms).

### Strain-specific isoform expression in mouse fibroblasts

To investigate mouse strain-specific isoform expression, we used all molecules with both an allele assigned and only a unique isoform assigned. We only considered genes for which we detected two or more isoforms and expression from both alleles. For each gene, we constructed a contingency table based on the counts of molecules assigned to each allele and isoform. Significance was tested was by using Chi-square test and the resulting p-values were corrected for the multiple testings using the Benjamini-Hochberg procedure. We further scrutinized the significant strain-isoform interactions (with an adjusted p-value < 0.05). For each significant gene, we performed thousand independent randomizations of allele and isoform labels of all molecules, and we computed the Chi-square test on each permutation, and we further required that the real p-value obtained were below 5% lowest p-values from the randomizations.

## References

1. Sandberg, R. Entering the era of single-cell transcriptomics in biology and medicine. Nat. Methods 11, 22–24 (2014).

2. Byrne, A. Nanopore long-read RNAseq reveals widespread transcriptional variation among the surface receptors of individual B cells. Nat. Commun. (2017).

3. Gupta, I. et al. Single-cell isoform RNA sequencing characterizes isoforms in thousands of cerebellar cells. Nat. Biotechnol. (2018) doi:10.1038/nbt.4259.

4. Mereu, E. et al. Benchmarking Single-Cell RNA Sequencing Protocols for Cell Atlas Projects. bioRxiv 630087 (2019) doi:10.1101/630087.

5. Ziegenhain, C. et al. Comparative Analysis of Single-Cell RNA Sequencing Methods. Mol. Cell 65, 631–643.e4 (2017).

6. Picelli, S. et al. Smart-seq2 for sensitive full-length transcriptome profiling in single cells. Nat. Methods 10, 1096–1098 (2013).

7. Deng, Q., Ramsköld, D., Reinius, B., & Sandberg, R. Single-cell RNA-seq reveals dynamic, random monoallelic gene expression in mammalian cells. Science 343, 193–196 (2014).

8. Bagnoli, J. W. et al. Sensitive and powerful single-cell RNA sequencing using mcSCRB-seq. Nat. Commun. 9, 2937 (2018).

9. Guo, J. U. & Bartel, D. P. RNA G-quadruplexes are globally unfolded in eukaryotic cells and depleted in bacteria. Science 353, (2016).

10. Ohtsubo, Y., Nagata, Y. & Tsuda, M. Compounds that enhance the tailing activity of Moloney murine leukemia virus reverse transcriptase. Sci. Rep. 7, 6520 (2017).

11. Cole, C., Byrne, A., Beaudin, A. E., Forsberg, E. C. & Vollmers, C. Tn5Prime, a Tn5 based 5’ capture method for single cell RNA-seq. Nucleic Acids Res. 46, e62 (2018).

12. Larsson, A. J. M. et al. Genomic encoding of transcriptional burst kinetics. Nature 565, 251–254 (2019).

13. Parekh, S., Ziegenhain, C., Vieth, B., Enard, W. & Hellmann, I. zUMIs - A fast and flexible pipeline to process RNA sequencing data with UMIs. GigaScience 7, (2018).

14. Reinius, B. et al. Analysis of allelic expression patterns in clonal somatic cells by single-cell RNA-seq. Nat. Genet. 48, 1430–1435 (2016).

15. Patro, R., Duggal, G., Love, M. I., Irizarry, R. A. & Kingsford, C. Salmon provides fast and bias-aware quantification of transcript expression. Nat. Methods 14, 417–419 (2017).

16. Martinez, N. M. & Lynch, K. W. Control of alternative splicing in immune responses: many regulators, many predictions, much still to learn. Immunol. Rev. 253, 216–236 (2013).

17. Wang, E. T. et al. Alternative isoform regulation in human tissue transcriptomes. Nature 456, 470–476 (2008).

18. Katz, Y., Wang, E. T., Airoldi, E. M. & Burge, C. B. Analysis and design of RNA sequencing experiments for identifying isoform regulation. Nat. Methods 7, 1009–1015 (2010).

19. Trapnell, C. et al. Differential analysis of gene regulation at transcript resolution with RNA-seq. Nat. Biotechnol. 31, 46–53 (2013).

20. Regev, A. et al. The Human Cell Atlas. eLife 6, (2017).

21. Scotti, M. M. & Swanson, M. S. RNA mis-splicing in disease. Nat. Rev. Genet. 17, 19–32 (2016).

## Supplementary Methods References

22. Picelli, S. et al. Full-length RNA-seq from single cells using Smart-seq2. Nat. Protoc. 9, 171–181 (2014).

23. Keane, T. M. et al. Mouse genomic variation and its effect on phenotypes and gene regulation. Nature 477, 289–294 (2011).

24. Lawrence, M. et al. Software for computing and annotating genomic ranges. PLoS Comput. Biol. 9, e1003118 (2013).

25. Stuart, T. et al. Comprehensive Integration of Single-Cell Data. Cell 177, 1888–1902.e21 (2019).

